# The effects of protein supplementation, fumagillin treatment, and colony management on the productivity and long-term survival of honey bee (*Apis mellifera*) colonies in Canada

**DOI:** 10.1101/2023.07.10.548417

**Authors:** Michael Peirson, Abdullah Ibrahim, Lynae P. Ovinge, Shelley E. Hoover, M. Marta Guarna, Andony Melathopoulos, Stephen F. Pernal

## Abstract

In this study, we intensively measured the longitudinal productivity and survival of 362 commercially managed honey bee colonies in Canada, over a two-year period. A full factorial experimental design was used, whereby two treatments were repeated across apiaries situated in three distinct geographic regions: Northern Alberta, Southern Alberta and Prince Edward Island, each having unique bee management strategies. In the protein supplemented treatment, colonies were continuously provided a commercial protein supplement containing 25% w/w pollen, in addition to any feed normally provided by beekeepers in that region. In the fumagillin treatment, colonies were treated with the label dose of Fumagilin-B^®^ each year during the fall. Our study shows that that neither treatment provided consistent benefits across all sites and dates. Fumagillin was associated with a large increase in honey production only at the Northern Alberta site, while protein supplementation produced an early season increase in brood production only at the Southern Alberta site. The protein supplement provided no long-lasting benefit at any site and was also associated with an increased risk of death and decreased colony size later in the study. Differences in colony survival and productivity among regions, and among colonies within beekeeping operations, were far larger than the effects of either treatment, suggesting that returns from extra feed supplements and fumagillin were highly contextually dependent. We conclude that use of fumagillin is safe and sometimes beneficial, but that beekeepers should only consider excess protein supplementation when natural forage is limiting.

## Introduction

In recent years, high rates of honey bee *(Apis mellifera* L.) colony death, queen loss, and poor colony growth have often been reported [1–3] and many have sought explanations [4]. Surveys of beekeepers often report that environmental and management factors such as poor queens, starvation, and small colony size are leading causes of death [1, 3]. Others have summarized major factors in terms of the four ’P’s - poor nutrition, pesticides, pathogens, and parasites [5, 6]. One parasite in particular, *Varroa destructor* Anderson & Trueman, is recognized as extremely important [7] and a variety of controls have been developed against it [8]. Some stressors, which include acute pesticide exposures, sustained pollen deprivation, and certain pathogens, produce identifiable symptoms. Others, such as covert viral infections [9] or nosema disease [10], can be largely invisible. As beekeepers cannot respond to problems they cannot identify or predict, many turn to preventative or prophylactic solutions, often having uncertain efficacy.

Protein supplements and the anti-fungal medication fumagillin are two of the most well-established practices used by beekeepers in Canada to promote colony health. Typically, protein supplements are supplied to colonies in spring before the first natural pollen is available [11], though they may also be fed in fall [12]), while fumagillin is routinely supplied in sugar syrup prior to winter [11].

Protein supplements are intended to compensate for a shortage or nutritional inadequacy in the pollen available to hives. Supplementation is a longstanding practice recommended to beekeepers when natural pollen may be limited [13]. Commercial protein supplements such as the one used in this study are often based on soy flour, brewer’s yeast, and sugar syrup, as recommended by Haydak [14–16], with a proportion of bee-collected pollen, although numerous other ingredients may be employed. Protein supplements for bees are popular and have been widely investigated in controlled conditions and short-term field studies, with inconsistent results [17–20].

Fumagillin is a treatment for the parasites *Vairimorpha apis*, and *Vairimorpha ceranae* (previously known as known as *Nosema apis* and *Nosema ceranae*) [21], which can shorten the lifespan of bees [17, 22–24]. *V. ceranae*, which is widespread in Canada [25], has been variously described as a severe [22] or a trivial [26] threat to honey bee colonies. Fumagillin has long been recommended for widespread prophylactic use [27], but some have questioned its effectiveness in controlling *V. ceranae* [28, 29]. In field trials, some authors have failed to detect colony-level economic benefits [30], but others have found increases in colony survival, size, and honey production. Nearly all have demonstrated reductions in Vairimorpha prevalence or spore load.

Between 2014 and 2016, we conducted a cohort study in which we attempted to evaluate the relative importance of nutrition, pesticides, parasites, pathogens, and general management in three Canadian beekeeping operations representing different regions and commercial beekeeping practices. Two treatments were applied in a factorial design: (1) protein supplements applied (or not applied) throughout most of the active beekeeping season, as a test of the hypothesis that nutritional deficiency was a cause of colony under-performance, and (2) a fall application of fumagillin applied (or not applied) to control nosema disease. In this, our first output from this large-scale investigation, we focus specifically on colony productivity and survival and examine differences in these measures across beekeeping regions. Our expectation was that both treatments would improve colony health and, when in used in combination, would have additive beneficial effects across all locations.

## Methods

### Colony management

In spring 2014, new honey bee colonies were established at sites in Canada which represented three common economic models of commercial beekeeping: hybrid canola seed pollination (*Brassica napus* L.) with limited honey production in Southern Alberta (SAB), honey production from fields of commodity canola in Northern Alberta (NAB), and lowbush blueberry (*Vaccinium angustifolium* Aiton) pollination with no surplus honey production in Prince Edward Island (PEI). The colonies in Northern Alberta belonged to Agriculture and Agri-Food Canada (AAFC) and were managed by personnel in the apiculture research program, while colonies at the other locations were owned and managed by commercial beekeepers. There were two or three apiary-level groups per region (Table S1), each with approximately 40 colonies, for initial totals per region of 117 (SAB), 123 (NAB) and 76 (PEI) colonies. All producers moved hives locally multiple times as part of normal management practice (Table S1), but during the bloom period of the principal crop (lowbush blueberries or canola), most PEI and all Alberta colonies were either within, or immediately next to, fields of that crop.

### Colony establishment

The colonies were established in late April 2014 by splitting local overwintered hives (SAB) or 1 kg packages of bees imported from New Zealand (NAB). In PEI, colonies were established from splits made at blueberry pollination sites in mid-June. The target size for splits at both locations was 8 frames of bees with 4 or 5 frames of brood. The standard management practices of the PEI beekeeper also included splitting colonies in late summer, and as such daughter colonies were also included in the study. Newly mated Carniolan queens (Kona Queen Company, HI, USA) were introduced to each colony established at the beginning of the study. Queens were marked with green paint on their thorax, and their wings were clipped to prevent swarming and aid in their identification. Both spontaneous and beekeeper-initiated re-queening were permitted, and new queens were marked and recorded when observed by project staff. In the case of re-queening by beekeepers, local stock was used.

### Routine management

Beekeepers used their own equipment and management procedures. All producers exclusively used Langstroth deep boxes. Colonies were in groups of four per pallet and were wintered outdoors in two Langstroth deep boxes. Both Alberta producers used an insulating wrap and cover on each pallet. The PEI beekeeper used only an insulating top cover. No producer used antibiotics for the control of brood diseases. All producers controlled *Varroa destructor* using Apivar® (amitraz) strips during the spring and fall (Alberta locations) or used Apivar® in summer and oxalic acid in winter (PEI).

## Experimental design

The trial was a fully crossed design with two main factors: 1) continuous protein supplementation, and 2) fall treatment for nosema disease. Table S2 details treatment and replicate numbers by site. Split colonies were always assigned to the parent colony’s treatment group.

### Protein supplement treatment

Beekeepers applied their normal early spring and fall feeding practices, which for Alberta producers included commercial protein supplements (Global Patties with 15% pollen; Global Patties, Airdrie, AB, Canada), to all colonies in spring. Subsequently, half the colonies at each apiary received no additional supplement, while the other half received protein patties prepared by Global Patties according to their standard recipe, but modified to include 25% pollen from Canadian prairie sources. The modified (w/w) recipe contained: sucrose syrup: 46%, distillers dried yeast: 15%, defatted soy flour: 14%, irradiated pollen: 25%. The supplement was provided continuously as consumed throughout the active beekeeping season, except during the canola bloom, when pollen was known to be abundant (Table S3).

Pollen used in the 25% protein supplement was trapped corbicular pollen from the Canadian prairies, and was irradiated (Iotron Technologies Corp, Port Coquitlam, BC, Canada) at 10 kGy before use. Major constituents of the pollen included *Brassica napus* L, *Trifolium* spp., *Melilotus* spp., and *Glycine max* (L) Merrill.

### Fumagillin treatment

Each fall, Fumagilin-B® (DIN: 02231180; Medivet Pharmaceuticals, High River, AB, Canada) was applied to half the colonies per apiary (Table S2). The method and timing of application varied among commercial beekeepers (Table S4) but was always administered at the fall label dose (200 mg fumagillin per colony). After treatment with fumagillin-treated syrup, or throughout the fall if fumagillin was delivered as a syrup drench, unmedicated 67% (w/w) sucrose syrup was supplied *ad libitum* until the colonies were wrapped for winter or the hives stopped accepting additional feed. Syrup was supplied to each colony individually in hive-top feeders. Untreated colonies received 67% sucrose without fumagillin continuously throughout the same period.

### Data collection procedures

#### Colony Assessments

Colonies were assessed at 11 time points which are nominally described as: May 2014, June 2014, August 2014, November 2014, April 2015, May 2015, June 2015, August 2015, November 2015, April 2016 and May 2016 (for precise dates, see Table S5). All assessments included a measurement of colony size, inspection for visible symptoms of disease, a search for the queen or indications of queen-rightness, and the collection of samples of adult bees from the broodnest.

#### Colony population measurements

Three methods of measuring colony size were employed. For May, June, and August assessments in 2014, bee and brood populations were estimated from a visual assessment. In May, June, and August 2015 and May 2016, a digital imaging method was employed. All adult bee assessments began at first light and ended with the start of bee flight in the morning, to ensure the entire hive population was measured. Sealed brood assessments were usually done later the same day. In November and April, due to the colder weather, colony populations were measured less invasively by estimating the size of the cluster of bees (Fig S1, in Supplement 1).

#### Visual assessments

For each frame within a hive, the area covered by bees or by sealed brood was estimated as described by Delaplane et al. [31]. Each measurement was performed by a single inspector. The inspector laid a grid of 1-inch (2.54 cm) squares over the frame and counted the number of squares filled with adult bees or sealed brood. The values from each frame were summed to yield brood and bee areas for the colony.

#### Conversion of area measurements to counts

Area measurements were converted to estimated numbers of bees or brood using the following conversion factors: 8.70 adult bees and 25.3 sealed brood cells per square inch (1.35 bees and 3.93 sealed brood cells per cm^2^). The conversion factors were estimated from 33 images of adult bees and 12 images of sealed brood taken in 2015 (Tables S6 & S7).

#### Digital images

Methods for counting bees with digital images have been previously described [32]. A specially constructed frame stand with an opaque back and translucent sides was used to ensure uniform light in the image. A Nikon D7100 camera with a Nikkor 105 mm Micro VR lens was set on a tripod approximately 2 m from the position of the frames, such that the edges of the frames would be just inside the boundaries of the image. Each side of each frame with bees or sealed brood was photographed. Frames were handled as quickly and gently as possible, working from the top of the hive to the bottom with minimal smoke. Unphotographed bees (bees on the box, bottom board, etc.) were visually estimated by area to the nearest equivalent of a quarter frame side; for this purpose, one side of a frame was assumed to hold 1200 bees [33].

#### Counting bees and brood from images

Images of adult bees or sealed brood for each colony were imported to an image analysis program, Honeybee Complete® 4.2 (WSC Scientific GmbH, Heidleberg, Germany). This software automatically identifies, labels, and counts each identified bee or sealed brood cell in images, but with some inherent errors. Each analyzed image was screened, and if necessary corrected, by a human observer. In the case of sealed brood, results were often close to 100% accuracy, and no adjustment was applied unless the number of false detections or failed detections in the image appeared to exceed 3% of the total (observer’s visual estimate).

Three methods of correction were used, as needed: 1) If individual bees or brood were concentrated in one part of the image, the area selection tool of the software was used to exclude other parts of the image from analysis; 2) For individual bee recognition errors, an adjustment was applied by manually selecting or deselecting using the software; or 3) For adult bee estimates, correction factors were applied to account for HoneyBee Complete’s systematic under-recognition of the true number of individuals in an image. Correction factors were calculated for images of typical quality in each combination of region and date (Table S8). Images of poorer quality were assigned a custom correction factor or were manually counted.

#### Cluster size measurements

For population assessments in late fall and early spring, we considered the temperature too cold to break the cluster and examine hives frame by frame. Instead, we estimated cluster size, which is the number of inter-frame spaces filled with bees to the nearest ¼ of a space, as viewed from above and below [34]. Observations from the top and bottom of each box were averaged to give the cluster score for the box; the value for the hive was the sum of the scores for the boxes.

#### Honey production and colony weights

Weight measurements used digital platform scales (model GBK 260A, Adam Equipment, Oxford, CT, USA or equivalent), or a hanging scale (Optima digital crane scale, Model WGB1483661, Global Industrial Canada, Toronto, ON, Canada).

Filled honey supers were weighed after removal from the colonies, and net honey contents were determined by subtracting either the empty weight of the super before placement on the colony (NAB) or an average empty weight (SAB). Net honey production for the hive was the sum for all honey supers collected during the production season (late June to early September). Colonies in PEI did not produce surplus honey that was harvested.

Feed stores before and after winter were determined by weighing colonies during the November and April inspections.

#### Queen and disease inspections, and sample collection

Inspections for queen and disease status, and samples of bees were either done the same day as the population size assessments, or the following day. Frames of bees were gently removed from the hive and examined in turn until either the queen had been found or all frames had been examined twice. While looking for the queen, inspectors noted any unusual looking adult bees – deformities, abnormal behaviours, or external parasites as well as any brood with signs of disease. Once the queen was found, the frame she was on was set gently outside the hive, and two samples of bees were taken from the remaining frames: 1/2 sample vial (∼150 bees) from a frame with honey and no brood, and 1 full sample vial (∼ 400 bees) from a frame with brood. In November and April, colonies in PEI were not inspected, and colonies in Alberta were inspected rapidly to avoid chilling bees and brood (pest and pathogen measurements from these samples will be the subject of upcoming manuscripts).

#### Pollen collection

We measured the effects of treatment and region on the quantity of pollen collected. In 2014, each research location used the pollen traps that were available locally, which for Southern Alberta were front entrance style traps (Propolis-etc, St. Mathieu de Beloeil, QC, Canada) and for Northern Alberta were Ontario Agricultural College-style traps [35], previously custom built. In 2015, all three locations used the front entrance style traps.

In 2014, pollen was collected once from each hive in Alberta and the traps were in place for a maximum of four days per colony during the canola bloom period. In 2015, in all three regions, a subset of 12 large colonies per apiary (three from each treatment group) was repeatedly sampled for 48-hour periods approximately every two weeks from mid-May until mid-September.

#### Colony viability

Data and samples were collected as long as any bees remained. If no bees remained, or the colony was queenless and broodless at the end of the study (May 2016), the colony was considered to have become non-viable at the midpoint between the last verifiably queenright date (when either the queen or all stages of worker brood were seen), and the next inspection date. Data from dates when the colony was nonviable were excluded from analyses.

### Statistical analysis

Analyses were conducted using R version 4.12, and R Studio build 372 [36, 37]. The project data file can be found in supplements 2&3, the R code in supplement 4, and the output file in supplement 5.

#### Cox proportional hazards models

Colony survival and queen survival were analyzed as Cox proportional hazards models, using the R Survival package, version 3.3-1 [38, 39]. Hazards were not proportional over time among regions; consequently, region was a stratified factor in the model. The split colonies at PEI could not be regarded as independent of their parents, so related colonies were grouped using a cluster command. Since fumagillin was not applied until the end of the first field season, some deaths among colonies assigned to the fumagillin treated group could not be attributed to fumagillin. To resolve this, the dataset was split into non-overlapping time periods for each colony as described by Therneau et al. [40].

#### Mixed models

Measures of colony population (adult bee counts, sealed brood counts, and cluster sizes), productivity (honey production, pollen collection), and syrup feed consumption (colony weight) were analyzed as dependent variables in linear mixed effects models using the R nlme package, version 3.1_153. For a comparison of linear mixed effects models with other approaches to handling repeated measures data, see Ma et al. [41]. Most dependent variables were modelled without transformation and with variances weighted by date or by date and region to reduce heteroskedasticity. The independent (fixed effects) variables under consideration were region, date, protein supplement treatment, and fumagillin treatment, and their interactions. Random intercepts and random slopes by colony, as well as several correlation structures were compared and the models with the best fit statistics were selected (S4 & S5). Final models were selected by successively removing non-significant higher order interactions until only interactions that were significant at P<0.05 or necessary to represent the experimental structure remained.

Some dependent variables (adult bees, brood, and honey production) were measured repeatedly both before and after the first fumagillin application in fall 2014; consequently, the main effect of “fumagillin” was not meaningful over all time points, however the interaction of fumagillin with date allowed interpretation of effects and was inherently implied by the project design. Therefore, both were retained in the models even if they were not significant at P<0.05.

#### Contrasts

Statistical significance (p) values for contrasts are shown without adjustment for multiple comparisons; instead, the significance threshold (p=0.05) was adjusted with a Bonferroni correction (S5). When the analysis of variance indicated that a factor and its interaction were both significant, we present contrasts for both. In such cases, the interactions were usually differences in the size of the treatment effect, not the direction, and as such the significant interaction does not exclude the more general result.

#### Adult bee count and sealed brood count models

There was no data for May 2014 at PEI and the practice there of splitting colonies in late summer complicated the analysis of colony populations. For simplicity, most statistics in the paper refer to colony sizes as observed (without adjusting for the effects of splitting). To clarify the influence of splitting, each statistical model was also tested on variants of the dataset that either excluded all split colonies, excluded colonies that were not split, or attributed all productivity of each daughter colony to the parent colony. These analyses are discussed only where the results differed from the main analysis. Because date was treated as a categorical fixed effect and the interaction of date and region was important, the lack of data from PEI in May 2014 meant that May 2014 data from the other sites had to be dropped from analysis. An alternative model was considered which would have allowed the whole dataset to be used by dividing the date term into a seasonal component (”month”) and a continuous component (”assessment”), but the results were similar to those presented.

#### Cluster size and colony weight models

The linear mixed models for cluster size and colony weight data included five fixed effects: “winter” (2014 or 2015), “month” (November or April), “region”, “patties”, and “fumagillin”. Colonies that died during winter (and therefore had no April measurement) were dropped from the November data for the same winter so that comparisons between spring and fall would reflect results from the same colonies. In addition, after a preliminary analysis we found that recently split colonies were smaller than other colonies, and the number of splits was not uniform among treatment groups, which resulted in a region-treatment interaction that was unrelated to the effects of treatment. Alternate models were used to study this problem (analyzing regions separately; analyzing split and non-split colonies separately; assigning all bees from the daughter split to the parent), but in the results presented, colonies split in August were excluded from the dataset for the following winter.

The linear models for the change in cluster size and change in weight during winter included four fixed effects: “winter” (2014 or 2015), “region”, “patties”, and “fumagillin”. We also considered the possibility that apparent treatment-related effects on the change in cluster size and colony weight might be functions of colony size at the beginning of winter. Therefore, cluster size in November, and its interactions, were added as predictors in the models for the change in weight and the change in cluster size.

#### Honey and pollen collection models

Honey production was modelled as a linear mixed model with fixed effects “year” (2014 or 2015), “region” (SAB or NAB), “patties”, and “fumagillin”.

Pollen collection data was square root transformed before analysis to reduce skewness. Different models were used for 2014 and 2015 data because the experimental procedure was different. In 2014, total pollen collected per colony was analyzed using a mixed model with three fixed factors: region (SAB or NAB), protein supplements (yes or no), and pollen collection date. Since each colony was tested only once, in this case the random factor was “apiary”.

In 2015, pollen was trapped repeatedly from a subset of colonies within each region. Fumagillin was not expected to affect pollen collection and was excluded from the model after a preliminary inspection of the data showed no evidence of an effect. The model contained three fixed effects: region (SAB, NAB, PEI), protein supplements (yes or no), and season (before canola bloom, during canola bloom, and after canola bloom). Season was used rather than date because the dates of testing were different for different colonies and sites, and to account for the fact that protein supplements were not applied during the canola bloom. Because repeated measurements were taken, “colony” was included as a random effect. Variances were weighted by date (not season) and region.

## Results

### Colony Survival

More than half of the colonies (194 of 362; 53.6%) died during the course of this two-year study. There were large differences in the numbers of deaths among the three beekeeping operations: 27% (32 of 117) of colonies in Southern Alberta, 47% (58 of 123) in Northern Alberta, and 85% (104 of 122) in PEI perished (Table 1; Fig 1A). Slightly fewer colonies died in the fumagillin treated groups, and slightly more in the protein supplemented groups, except in Northern Alberta, where most colonies in the protein supplemented groups died. The reduction in risk associated with fumagillin was not statistically significant (χ^2^ = 1.284; df=1; p= 0.257; hazard ratio: 0.84 ± 0.13; z = -1.13; p=0.26). The protein supplement significantly increased the death rate, both as a main effect (χ^2^ = 5.325; df=1; p= 0.021) and as an interaction with region (χ^2^ = 6.493; df=2; p= 0.039). In Southern Alberta and PEI, however, the increased death rate was not significant (SAB: Fig 1B; hazard ratio: 1.02 ± 0.36; z= 0.058; p=0.95; PEI: Fig 1d: hazard ratio: 1.14 ± 0.21; z= 0.72; p=0.47). Protein supplements were associated with a significantly higher death rate in Northern Alberta (Fig 1C; hazard ratio: 2.45 ± 0.65; z= 3.36; p<0.001)

**Fig 1.**
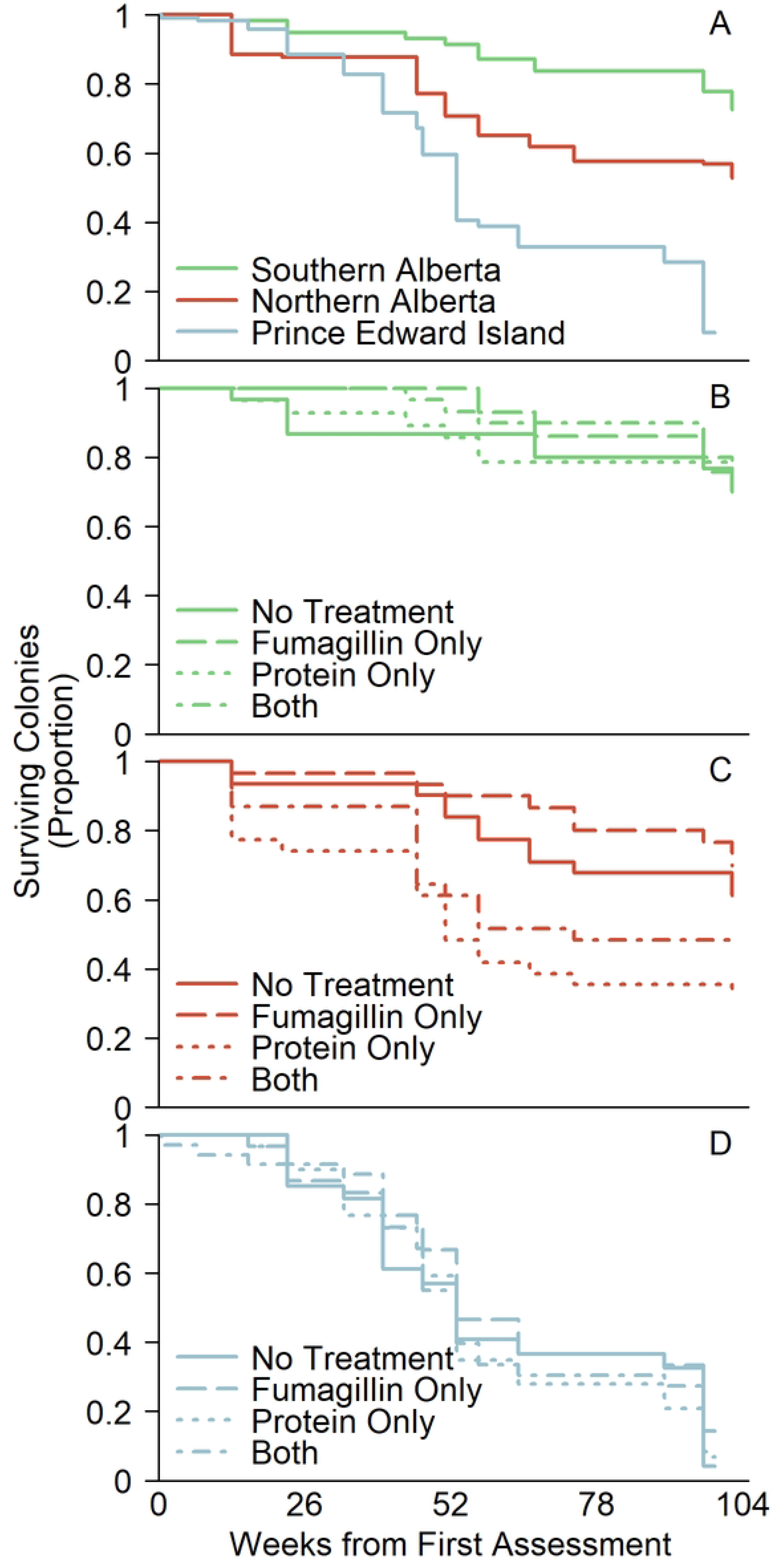
Colony survival curves. (A) Colony survival by region. (B) Effect of treatment on colony survival in Southern Alberta. (C) Effect of treatment on colony survival in Northern Alberta. (D) Effect of treatment on colony survival in PEI.

**Table 1.** Colony and queen survival to the end of the experiment.^a^

| Region & treatment | Total colonies | Colony deaths | Surviving colonies<sup>c</sup> | Surviving original queens<sup>c</sup> | Successful queen replacements<sup>d</sup> |  |
| --- | --- | --- | --- | --- | --- | --- |
|  | (n) | (%) | (n) | (n) | Total | After fumagillin treatment<sup>e</sup> |
| Southern Alberta | 117 | 27 | 85 | 47 | 50 | 40 |
| • Control | 30 | 30 | 21 | 13 | 11 | 8 |
| • Fumagillin | 29 | 24 | 22 | 15 | 12 | 9 |
| • Protein | 28 | 29 | 20 | 9 | 12 | 10 |
| • Fumagillin & | 30 | 27 | 22 | 10 | 15 | 13 |
| Northern Alberta | 123 | 47 | 65 | 14 | 92 | 36 |
| • Control | 31 | 39 | 19 | 5 | 21 | 10 |
| • Fumagillin | 30 | 30 | 21 | 5 | 28 | 12 |
| • Protein | 31 | 68 | 10 | 1 | 21 | 7 |
| • Fumagillin & | 31 | 52 | 15 | 3 | 22 | 7 |
| Prince Edward Island<sup>b</sup> | 110 | 85 | 13 | 3 | 37 | 32 |
| • Control | 24 | 96 | 1 | 0 | 8 | 8 |
| • Fumagillin | 28 | 79 | 6 | 2 | 9 | 9 |
| • Protein | 26 | 88 | 3 | 1 | 11 | 8 |
| • Fumagillin & | 32 | 91 | 3 | 0 | 9 | 7 |

<sup>a</sup> Data shown reflect colony assignment to treatment group. Some colony deaths occurred before the first fumagillin application, and thus do not reflect the effectiveness of that treatment.
<sup>b</sup> Data for PEI includes original colonies and splits made in 2014 but not the 12 additional splits produced in August 2015.
<sup>c</sup> At the May 2016 inspection date.
<sup>d</sup> Includes both natural and beekeeper-initiated queen replacements. Excludes queen failures associated with colony death. Some colonies experienced multiple queen events. These numbers represent known queen replacements.
<sup>e</sup> The first fumagillin application did not occur until September 2014. Queen deaths after fumagillin treatment are shown separately because fumagillin has previously been recommended for preventing queen loss [26].

Significantly more colony deaths occurred during the winters (98 deaths between the November and May inspections), than during the summers (84 deaths between May and November; χ^2^ = 4.528, df = 1, p = 0.033). The high colony loss rate at PEI was partly associated with colony splitting and feeding practices employed by the cooperating beekeeper (Table 2). In 2014, colonies that were split were less likely to die before winter than colonies that were not split, which probably reflects that the beekeeper chose to split successful colonies (χ^2^ = 4.62, df = 1, p = 0.032). These splits were *more* likely to die during winter (χ^2^ = 6.70, df = 1, p-value = 0.010).

**Table 2.**
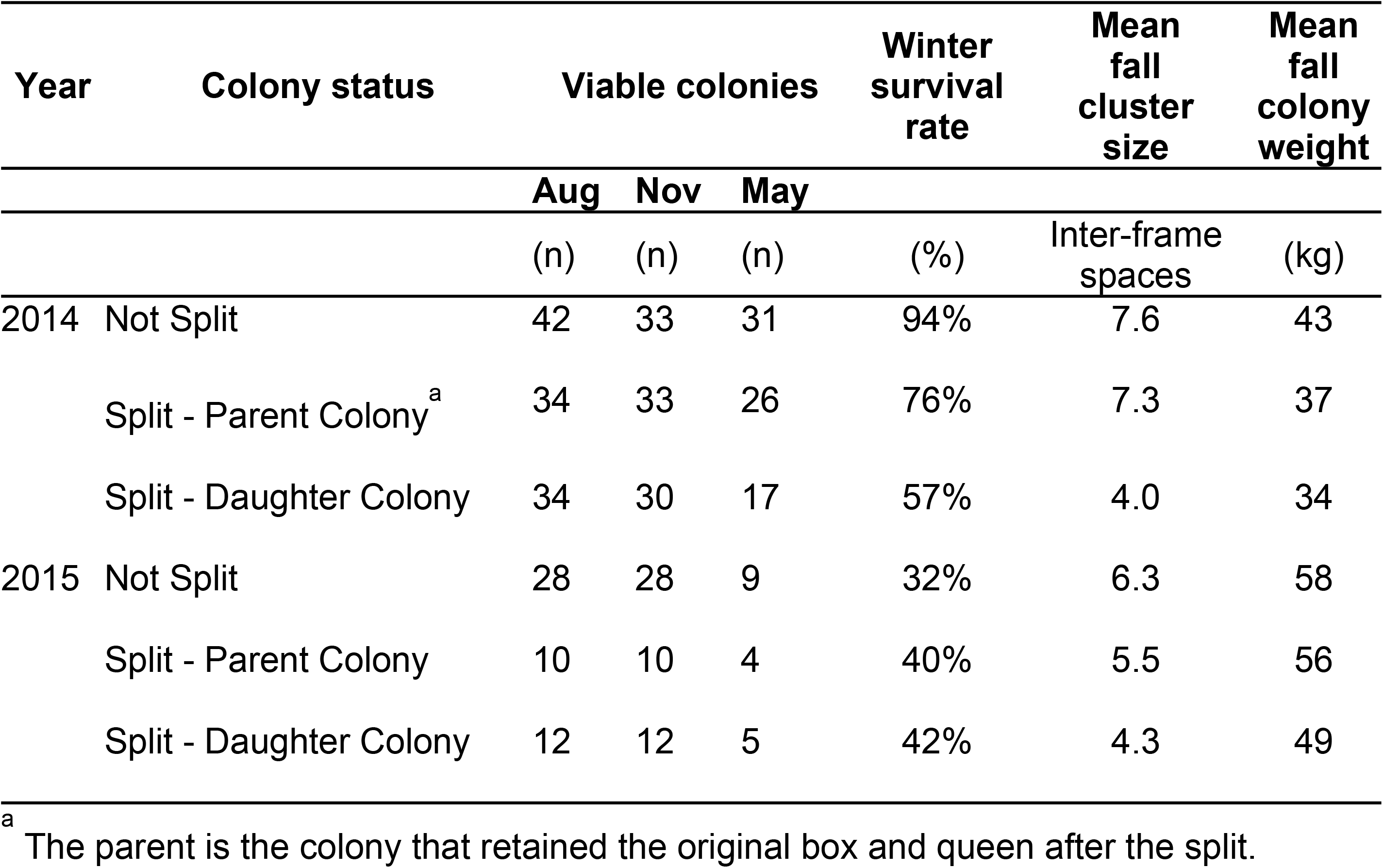
Characteristics of split and non-split colonies at PEI.

| Year | Colony status | Viable colonies |  |  | Winter survival rate | Mean fall cluster size | Mean fall colony weight |
| --- | --- | --- | --- | --- | --- | --- | --- |
|  |  | Aug | Nov | May |  |  |  |
|  |  | (n) | (n) | (n) | (%) | Inter-frame spaces | (kg) |
| 2014 | Not Split | 42 | 33 | 31 | 94% | 7.6 | 43 |
|  | Split - Parent Colony<sup>a</sup> | 34 | 33 | 26 | 76% | 7.3 | 37 |
|  | Split - Daughter Colony | 34 | 30 | 17 | 57% | 4.0 | 34 |
| 2015 | Not Split | 28 | 28 | 9 | 32% | 6.3 | 58 |
|  | Split - Parent Colony | 10 | 10 | 4 | 40% | 5.5 | 56 |
|  | Split - Daughter Colony | 12 | 12 | 5 | 42% | 4.3 | 49 |

<sup>a</sup> The parent is the colony that retained the original box and queen after the split.

### Queen Survival

Twenty percent of the original queens (63 of 316) were still alive at the end of the two-year study. Thirty-eight percent of surviving colonies were still headed by their first queen (64 of 168, taking into account original colonies and splits made in 2014, but not splits made in 2015; Table 1). Most of the surviving original queens (47; 73%) were in Southern Alberta. Fewer colonies in the protein supplemented groups retained their original queen (24 of 73 protein supplemented colonies versus 39 of 90 unsupplemented). The group that received fumagillin without additional protein had the greatest percentage of surviving original queens (45%; 22 of 49 surviving colonies still had their original queen) as well as having had the greatest percentage of surviving colonies. However, these differences in queen survival frequency were not statistically significant (χ^2^ = 1.87, df = 3, p = 0.60).

The timing of queen events, and the number of queen events differed among regions. At least 92 successful queen replacements occurred in Northern Alberta; nearly all were natural supersedures or failed swarms, and most of these occurred during the first summer. Some colonies experienced multiple queen changes. A Cox Proportional Hazards test for queen survival, using only queen events that were not associated with colony death, did not reveal significant differences associated with the treatments, but the effect estimates resembled those for the Cox model of colony survival. Queens in protein supplemented colonies tended to have shorter survival times while those in fumagillin treated colonies had longer survival times (S5). Many colony deaths were known to have been preceded by a queen loss, but we could not distinguish such deaths in every case. Therefore, we also considered a combined model in which all colony deaths were treated as a type of queen event (representing the case where queen replacement failed). In that model, protein supplements significantly reduced queen survival times regardless of region (ANOVA: χ^2^ = 6.619; df=1; p= 0.010; effect estimate: hazard ratio = 1.26 `±` 0.12; z=2.52; p=0.012). The effect of fumagillin still was not significant (ANOVA: χ^2^ = 1.463; df=1; p= 0.227; effect estimate: hazard ratio = 0.87 `±` 0.10; z=-1.21; p=0.23).

### Colony population measurements

#### Effects of region and date

For each dependent measure of colony population, differences related to region and date were highly statistically significant (p<0.001) and greatly outweighed the effects of treatments. For simplicity, these comparisons are not reported in subsections below, but may be found in Supplement 5.

The colonies in Southern Alberta, which had been started as nucleus hives, were initially much larger (9,200 ± 300 adult bees per colony at the May 2014 inspection; Fig 2A) than those in Northern Alberta, which had been started from packages (3,900 ± 130 adult bees; Fig 2b). Nevertheless, the Northern colonies already had more brood on that date (Figs 3A & B) and grew more quickly, reaching a peak of 25,000 ± 920 adult bees per colony in August 2014.

**Fig 2.**
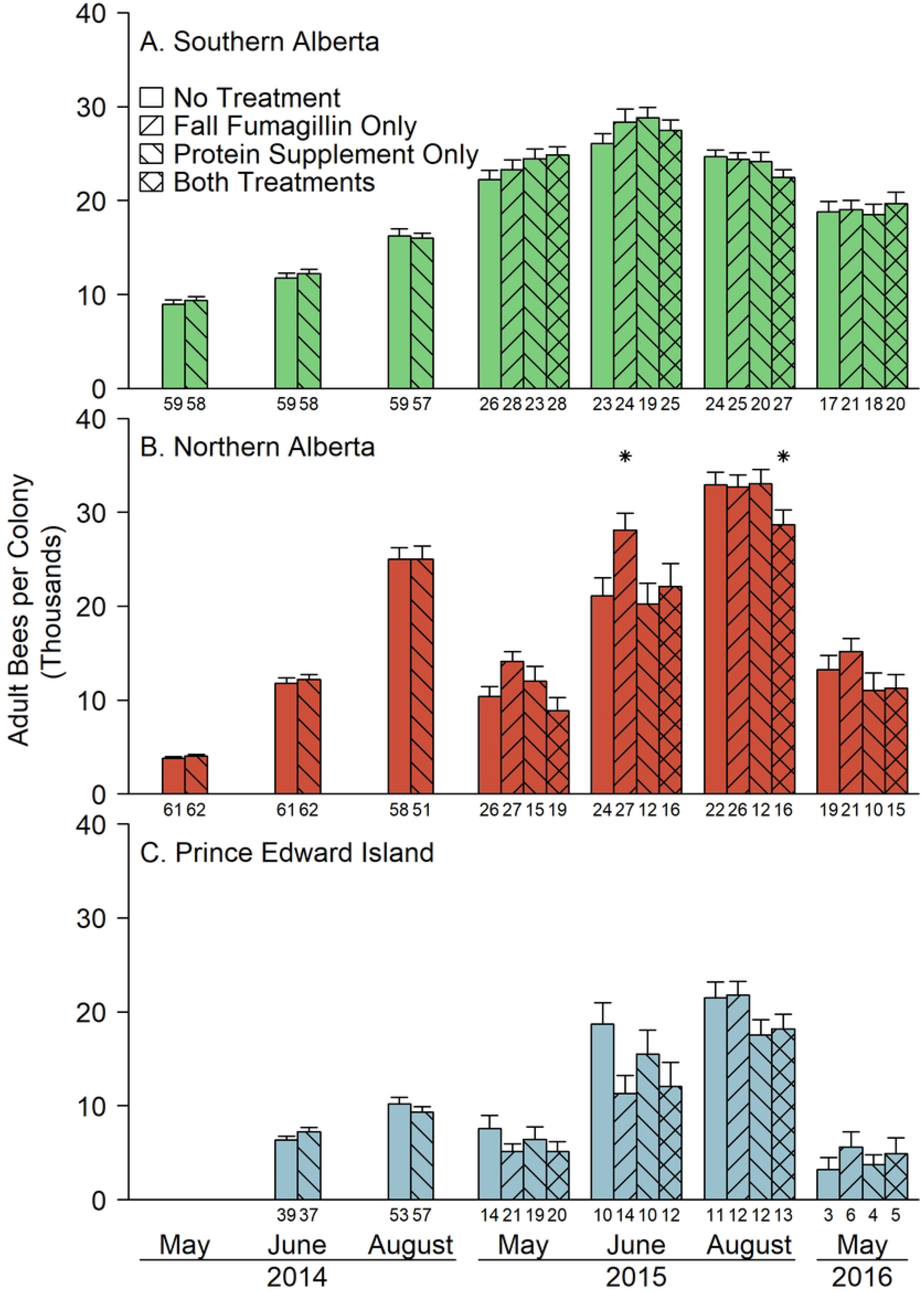
Colony size measured as adult population per colony (mean ± SE). Stars (*) above a column indicate that the treatment group was significantly different from the untreated control group in contrasts within region and date (p<0.05, Bonferroni adjusted; see Supplement 5). Numbers below a column indicate the number of viable colonies in the treatment group. Fumagillin was first applied in the fall of 2014; consequently, only two columns (untreated and protein supplemented) are shown on the earlier dates; and these columns include colonies subsequently treated with fumagillin.

**Fig 3.**
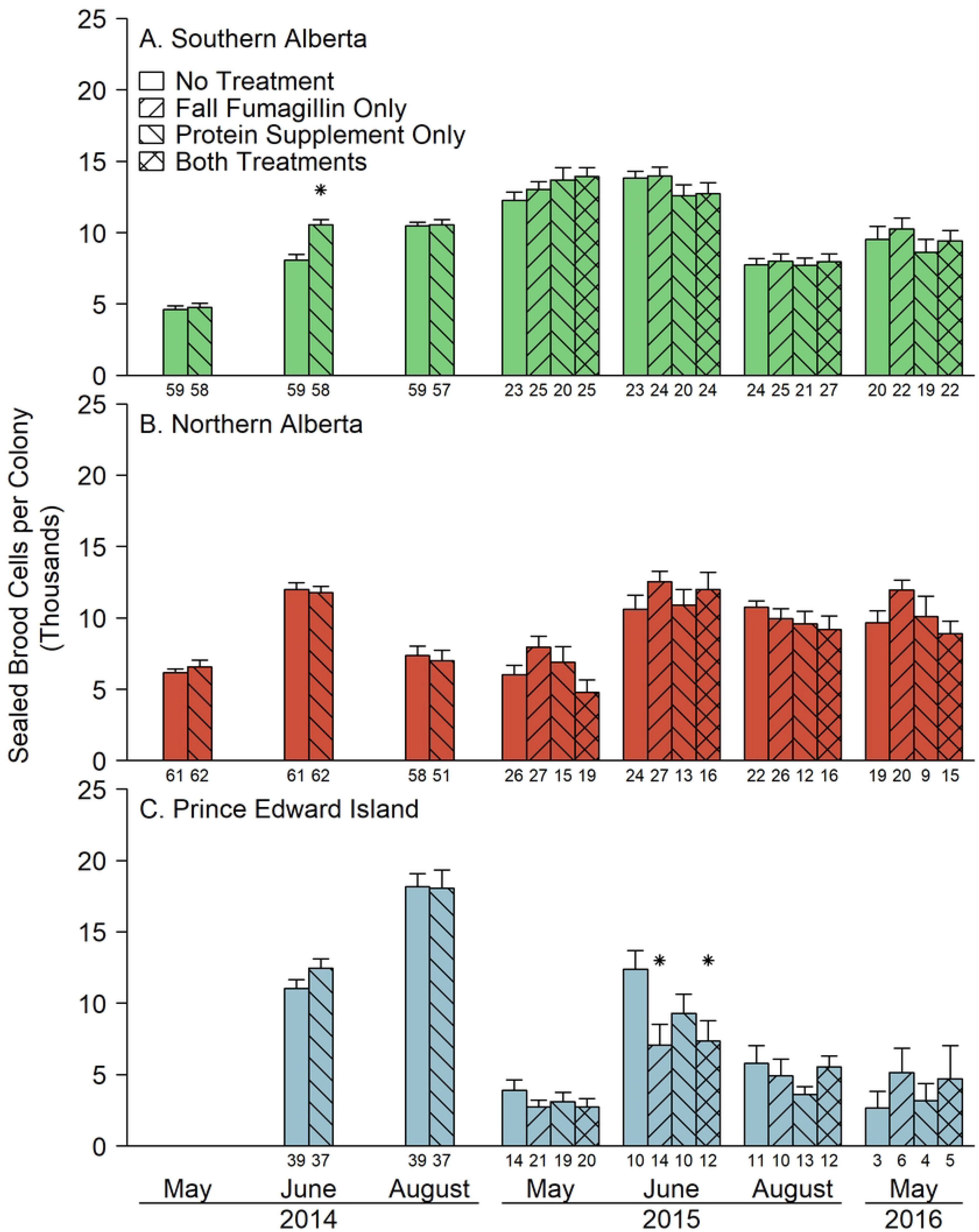
Colony size measured as the number of sealed brood cells per colony (mean ± SE). Stars (*) above a column indicate that the treatment group was significantly different from the untreated control group in contrasts within region and date (p<0.05, Bonferroni adjusted; see Supplement 5). Numbers below a column indicate the number of viable colonies in the treatment group. Fumagillin was first applied in the fall of 2014; consequently, only two columns (untreated and protein supplemented) are shown on the earlier dates; and these columns include colonies subsequently treated with fumagillin.

The initial (June 2014) adult bee population in PEI was low (6,800 ± 290 bees per hive), but these colonies had nearly as much sealed brood (Fig 3C) as the Northern Alberta colonies. PEI colonies were split in late summer, following the beekeeper’s standard management practice. Without splitting, the average colony would have contained 14,100 ± 600 bees in August 2014 (not shown), but as observed, populations were only 9,700 ± 450 adult bees (Fig 2C). All the sealed brood was left with the original parent colonies which, consequently, had the most brood of any site or date (18,100 ± 770 sealed brood cells; Fig 3C); splits contained only open brood at the time of inspection (not shown).

Changes in colony populations during winter (Fig 4) corresponded to the length and timing of the season. Cluster measurements (Fig 4 A & B) were not equivalent between Alberta regions because temperatures were warmer in the south and bees did not cluster tightly. In PEI, the onset and the end of cold weather were both much later than in Alberta, and consequently cluster measurements were also much later. After winter, colonies in Southern Alberta were by far the largest and colonies at PEI were the smallest, while Northern Alberta colonies, which experienced the longest period of cold weather, also experienced the largest reduction in size (Fig 4).

**Fig 4.**
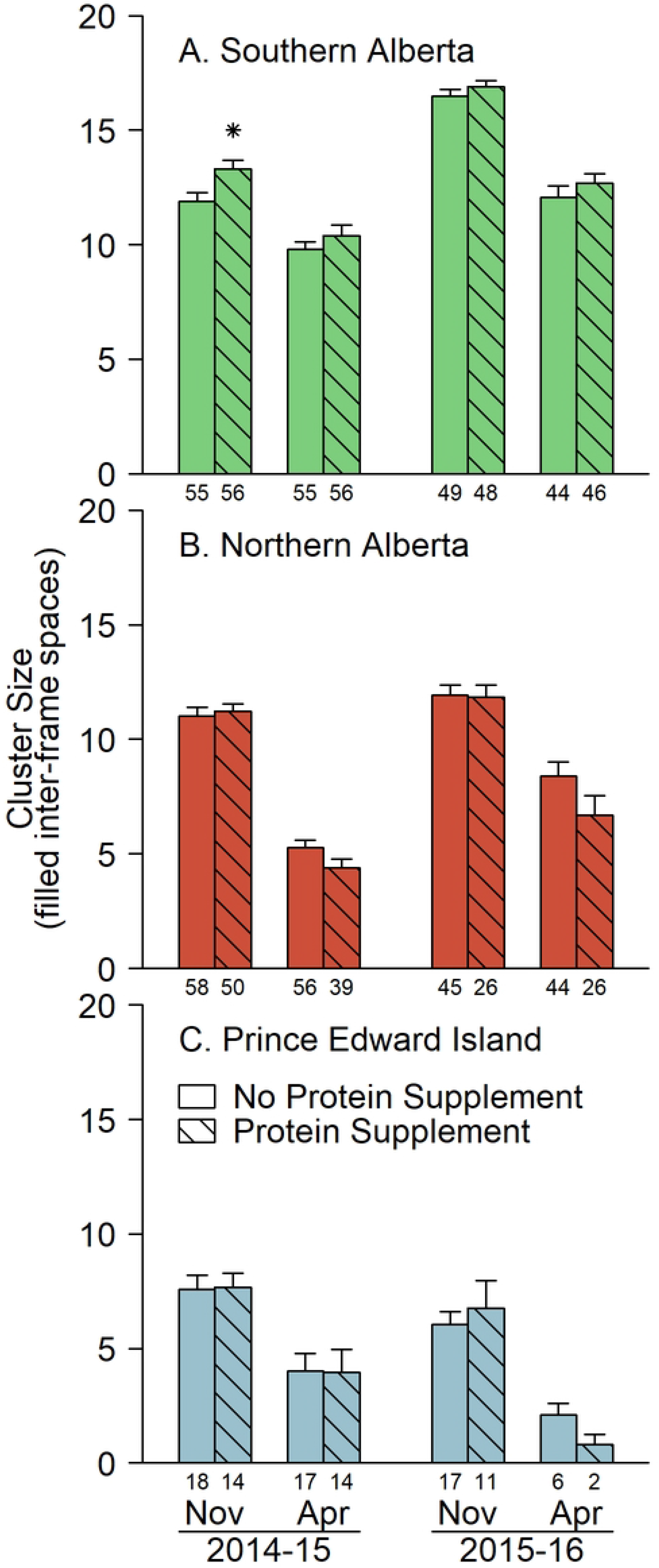
Colony size measured as the number of inter-frame spaces filled with clustering bees (mean ± SE). Stars (*) above a column indicate that the protein supplemented group was significantly different from the unsupplemented group in contrasts within region and date (p<0.05, Bonferroni adjusted; see Supplement 5). Numbers below a column indicate the number of viable colonies in the treatment group. Fumagillin did not affect cluster sizes; therefore, columns include both fumagillin treated and untreated colonies.

By the mid-May 2015 inspection, colonies in the south had produced several generations of brood, but many hives in PEI had not yet produced emerging bees. Regional patterns of colony growth in the second year mirrored the first except that the surviving colonies were larger (Figs 2-4).

#### Effects of splitting on colony size

The late summer splits at PEI created a complicating factor. Though the experimental protocol required nucleus colonies be created in spring 2014, the beekeeper also split colonies in late summer, in keeping with their standard management. Table 2 shows the numbers of split and un-split colonies, their rates of survival to spring, and average weights and cluster sizes before winter. The splits were a beekeeper decision and consequently, the treatment groups were not equally affected. Protein supplemented colonies were split more frequently (a marginally significant effect when both years are combined: χ^2^ = 3.2, df=1, p = 0.07). The decision to split colonies would presumably have been based on perceived colony size, but our measurements do not support the view that protein supplemented colonies were larger before splitting. For example, in August 2014, if we attribute all adult bees in the splits to the parent colony, adult populations in PEI before splitting were 14,300 ± 820 in the protein supplemented group and 13,900 ± 860 in the unsupplemented group, which is not a significant difference (two-sided t test: t = 0.384, df = 73.9, p-value = 0.70).

In both years, colonies that had been split were significantly smaller in November than colonies that had not been split (2014: t=3.40, df=70.2, p=0.001; 2015: t=2.06, df=46.4, p=0.045), and among split colonies, daughters were significantly smaller than parents in 2014 (2014: t=5.62, df = 40.1, p<0.001; 2015: t=1.37, df=18.4, p=0.18). Colonies that were split weighed less before winter than colonies that were not split, and daughter colonies from splits weighed less before winter than parent colonies (2014, not split versus split: t=. 5.23, df=66.2, p<0.001; 2015, not split versus split: t=1.71, df=32.12, p=0.097; 2014, daughter versus parent: t=2.02, df=58.0, p=0.049; 2015, daughter versus parent: t=1.71, df=32.1, p=0.31).

The average surviving PEI colony weighed just over 25 kg in spring 2015, which is similar to the weight of a two-box Langstroth hive that does not contain any honey as feed. Thus, it is probable that starvation may have contributed to the high death rate and small colony size in PEI, especially among split colonies.

#### Effects of treatments

Protein supplements did not increase the adult bee count in any region at any measurement date (Fig 2). On the contrary, protein supplements reduced the number of adult bees later in the experiment (interaction of protein supplements and date: F = 2.99; df=5,1115; p=0.011). Averaging the last five adult bee count dates across all three regions, supplemented colonies had 1,020 ± 410 fewer adult bees per colony (t=-2.48, df=350, p=0.014). This difference was not statistically significant on any individual date except August 2015 (-2,150 ± 650 adult bees; t=-3.31, df=350, p=0.001). By region (Fig 5A), the negative effect of protein supplements on adult bee population was significant only in Northern Alberta. Protein supplemented colonies in Northern Alberta had fewer adult bees across all dates after June 2014 (-1,760 ± 680 adult bees per colony; t=-2.584, df=350, p=0.010) and in August 2015 alone (-2,890 ± 850; t=-3.386, df=350, p<0.001).

**Fig 5.**
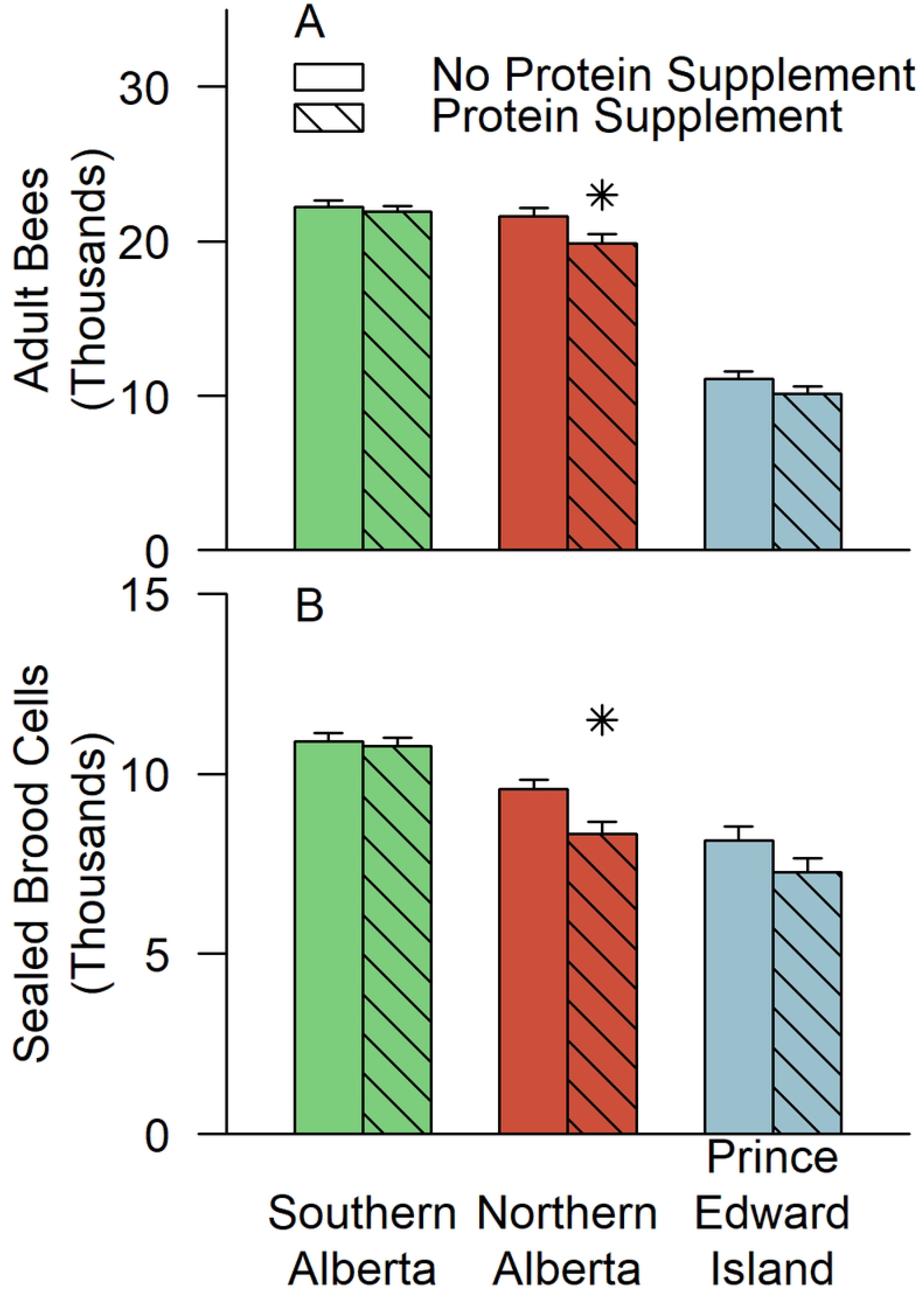
The effect of protein supplements on colony size late in the experiment. Data shown are estimated marginal means for (A) adult bee population and (B) sealed brood population averaged over the levels of the fumagillin treatment and the last five assessment dates (i.e., August, 2014, May 2015, June 2015, August 2015, and May 2016). Stars (*) above a column indicate that the supplemented group was significantly different from the unsupplemented group within region (p<0.05, Bonferroni adjusted; see Supplement 5).

Early in the experiment, in June 2014, protein supplemented colonies produced more sealed brood (Fig 3) than standard colonies across all regions combined (+1330 ± 400 sealed brood cells per colony; t = 3.34, df = 333, p < 0.001) and in Southern Alberta alone, this difference was substantial (difference by contrast: +1,960 ± 460 sealed brood cells per colony; t=4.30, df=333, p<0.001). On all other dates, however, protein supplemented colonies had, on average, less brood (protein * date: F = 5.37, df = 5,1091, p<0.001). Combining the last 5 brood inspection dates of the study, protein supplemented colonies had less brood in all regions (-750 ± 250 sealed brood cells; t=-2.99; df=333, p=0.003) and particularly in Northern Alberta (-1,250 ± 410 sealed brood cells; t=-3.02, df=333, p=0.003; Fig 5b).

Fumagillin treatments were applied each fall (between the August and November inspections) in feed or as a drench (Table S4). The three-way interaction of region, date, and fumagillin was significant in models for adult bees and sealed brood (Adult Bees: F=1.96, df= 10,1115, p=0.034; Sealed Brood: F=2.01, df=10,1091, p=0.029). Significant effects were detected by contrasts only in June 2015, when fumagillin-treated colonies in Northern Alberta contained more bees and brood than untreated colonies, while treated colonies in PEI contained fewer bees and brood than untreated colonies (Figs. 2 & 3). Neither effect reached significance in contrasts within region and date when averaged across levels of the protein supplement treatment, after accounting for multiple comparisons. However, in Northern Alberta, among colonies that were not receiving extra protein supplements, fumagillin was associated with significantly larger adult bee populations (Fig 2b). The surprising negative effect of fumagillin on colony populations in PEI in June 2015 (Figs 2c and 3c) arose entirely among colonies which had been split in August 2014. There were more splits in the fumagillin treated group and the fumagillin-treated split colonies were smaller both before and after winter. Splitting (Table 2) and fumagillin supplied in sugar syrup (see “Colony weight”, below) both reduced the amount of feed stored by hives prior to winter, which made these colonies particularly vulnerable, although death rates were similar. Two-thirds of fumagillin-treated splits weighed less than 40 kg prior to winter.

Cluster sizes were larger in the second winter than in the first (Fig 4). Fumagillin did not affect cluster sizes (F=0.358, df = 1,235, p=0.550). In contrasts within region and date, the only significant treatment effect was in Southern Alberta (Fig 4a). There, in November 2014, protein supplemented colonies were 1.45 ± 0.40 inter-frame spaces larger (t=3.649, df = 235, p < 0.001) than standard colonies. However, the four-way interaction of protein, region, year, and month was not significant in the analysis of variance.

In the first year, protein supplemented and standard colonies were the same size (averaged across levels of month, region, and fumagillin treatment; Figure 6A), but the supplemented colonies were 0.87 ± 0.33 inter-frame spaces of bees smaller in the second winter (t = -2.65, df = 235, p = 0.009). Additionally (Fig 6B), supplemented and standard colonies were similarly sized in fall (averaged across levels of year, region, and fumagillin treatment) but by spring, protein supplemented colonies were 0.66 ± 0.32 inter-frame spaces of bees smaller (t=-2.07, df = 235, p=0.039). The change in cluster size during winter also approached significance. Colonies that had received protein supplements declined 0.66 ± 0.34 inter-frame spaces of bees more than standard colonies (t = 1.93, df = 235, p = 0.055).

**Fig 6.**
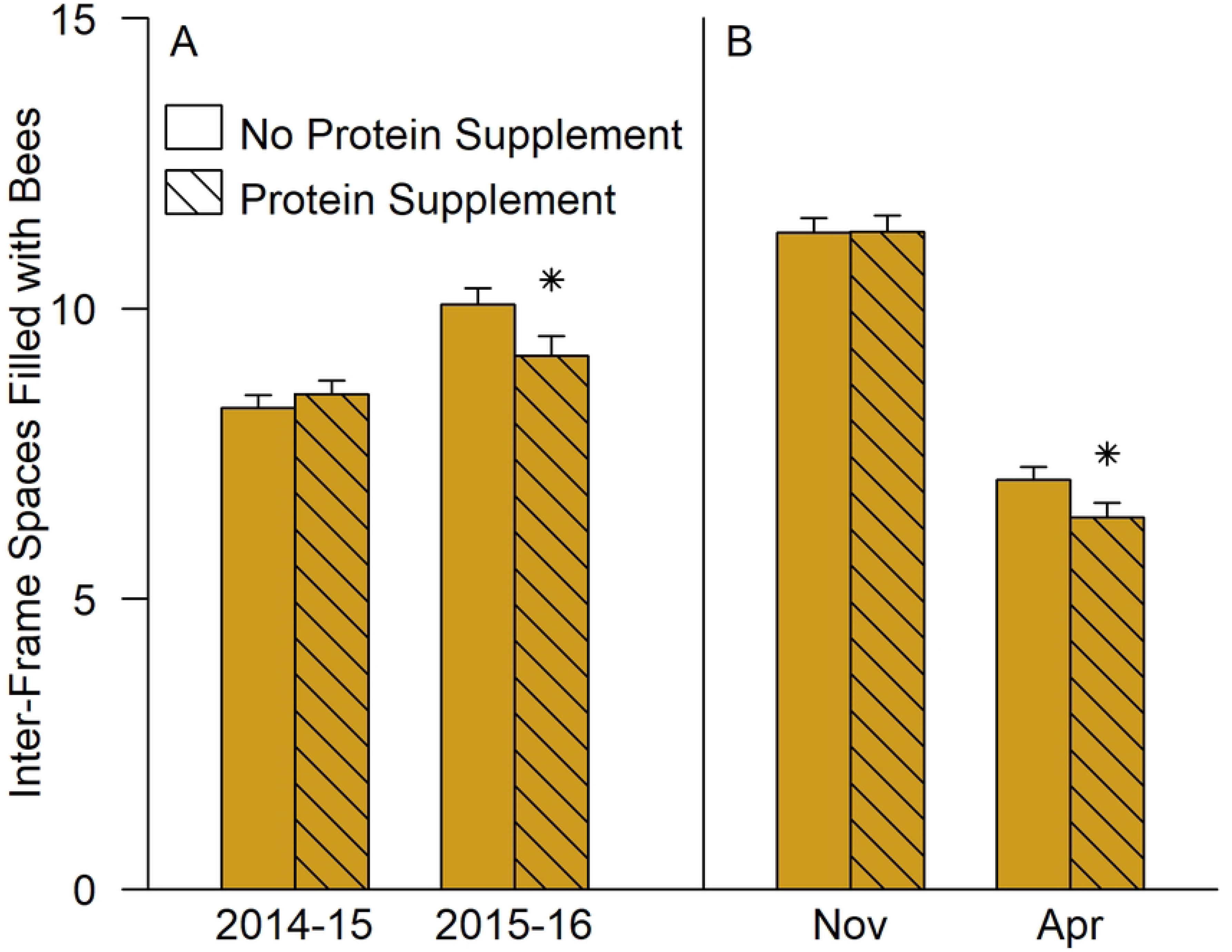
The model effect of protein supplements on cluster size (mean ± SE). Estimated marginal means are shown averaged across the levels of region and fumagillin. (A) Interaction of protein supplements and year (averaged across month): (B) Interaction of protein supplement and month (averaged across year). Stars above a supplemented column indicate a significant difference from the corresponding unsupplemented group (p<0.05, Bonferroni adjusted; see Supplement 5).

A significant two-way interaction between region and protein supplements was also detected (F=4.00, df = 2, 235, p=0.020). No within-region contrast quite reached significance, but supplemented colonies were relatively large in Southern Alberta and smaller at the other two locations (Fig 4).

#### Colony weight

Colony weights were measured at the November and April inspection dates as an indicator of feed storage and consumption. Colony weights were greater in fall, and in the second year of the study, and varied among regions according to the feeding practices of the beekeeper (Table 3). Colonies in PEI weighed far less than either Alberta site in the first winter, and many were near starvation or had starved by spring 2015. In PEI, colonies that were split were lighter than colonies that were not split, and daughter colonies were lighter than parent colonies (Table 2).

**Table 3.** Weights of honey bee colonies before and after winter.

| Region | Winter | Fall weight<sup>a</sup> | Spring weight | Average feed consumption |  |
| --- | --- | --- | --- | --- | --- |
|  |  | kg | kg | kg/winter | g/day |
| SAB<sup>b</sup> | 2014-15 | 56.7 ± 0.7<sup>c</sup> | 40.6 ± 0.6 | 16.1 | 109 |
| SAB | 2015-16 | 56.6 ± 0.4 | 38.3 ± 0.5 | 18.3 | 114 |
| NAB | 2014-15 | 56.4 ± 0.5 | 38.1 ± 0.5 | 18.3 | 108 |
| NAB | 2015-16 | 64.5 ± 0.7 | 40.7 ± 0.6 | 23.8 | 123 |
| PEI | 2014-15 | 39.6 ± 0.8 | 25.7 ± 0.5 | 13.9 | 86 |
| PEI | 2015-16 | 57.7 ± 2.6 | 37.6 ± 2.1 | 20.1 | 137 |

<sup>a</sup> Fall and spring data both exclude colonies that were not viable at the April weight measurement
<sup>b</sup> Abbreviations: SAB: Southern Alberta; NAB: Northern Alberta; PEI Prince Edward Island.
<sup>c</sup> Mean ± SE

Protein supplemented colonies were 2.01 kg ± 0.57 kg (t=3.513, df = 235, p<0.001) heavier than standard colonies before winter, but not after (Fig 7), indicating that more feed was stored and consumed. Fumagillin, in contrast, reduced average colony weight (Fig 8) (F = 23.77, df= 2, 235, p < 0.001) but its effect size ranged from zero to 5 kg per colony depending on region (F = 5.735, df= 2, 235, p = 0.004) and there was a three-way interaction with year and region (F = 10.71, df=2, 551, p < 0.001). The effect of fumagillin did not depend on month, which indicates that fumagillin reduced feed acceptance and storage, but not consumption during winter. One of the six within-region-and-date contrasts was statistically significant (Southern Alberta, first winter; t = 4.875, df = 235, p < 0.001; Fig 8).

**Fig 7.**
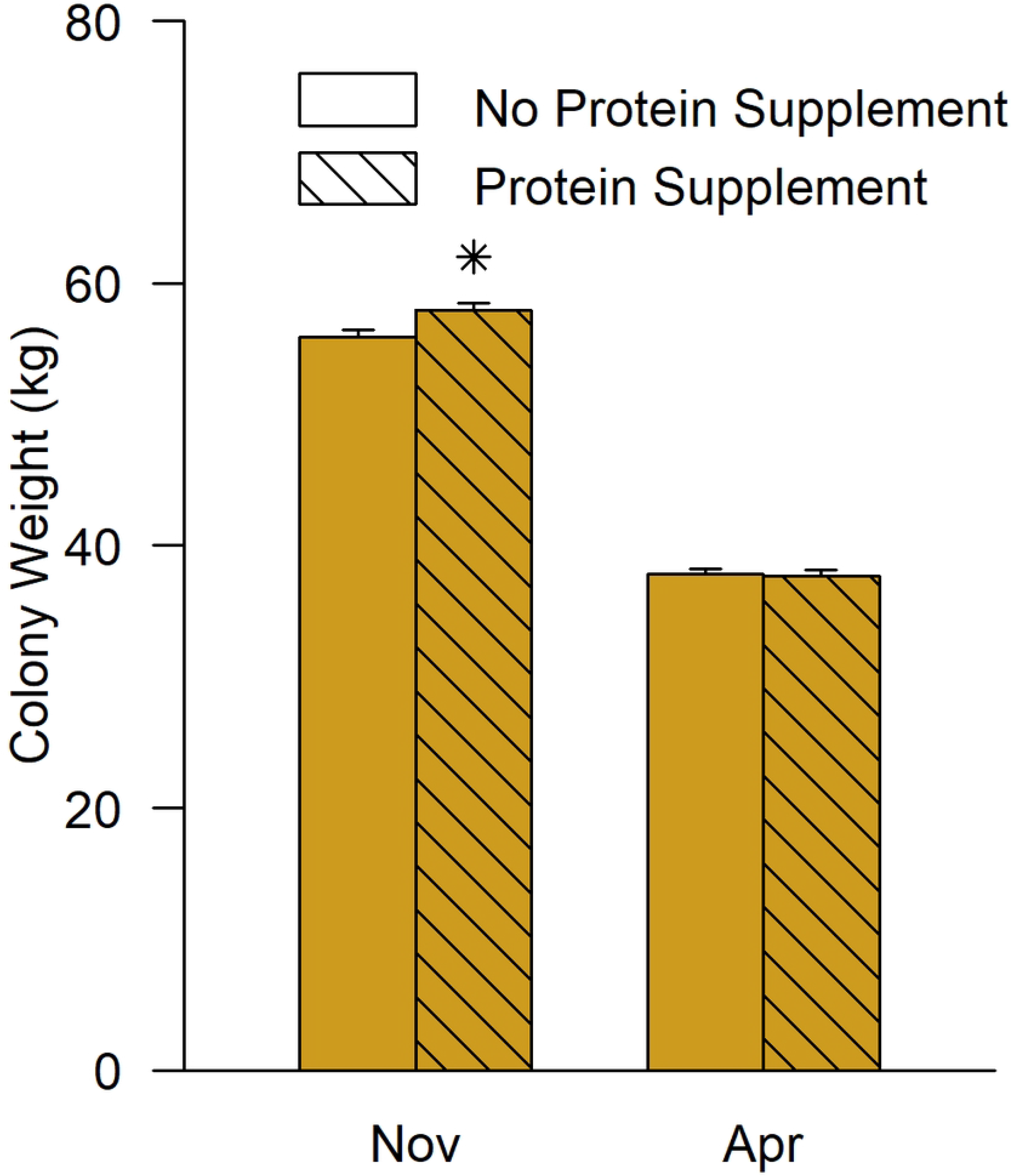
The model effect of protein supplements on colony weight (mean ± SE). Estimated marginal means are shown averaged across region, year, and levels of the fumagillin treatment. Stars above a column indicate that the supplemented group was significantly different from the unsupplemented group (p<0.05, Bonferroni adjusted; see Supplement 5).

**Fig 8.**
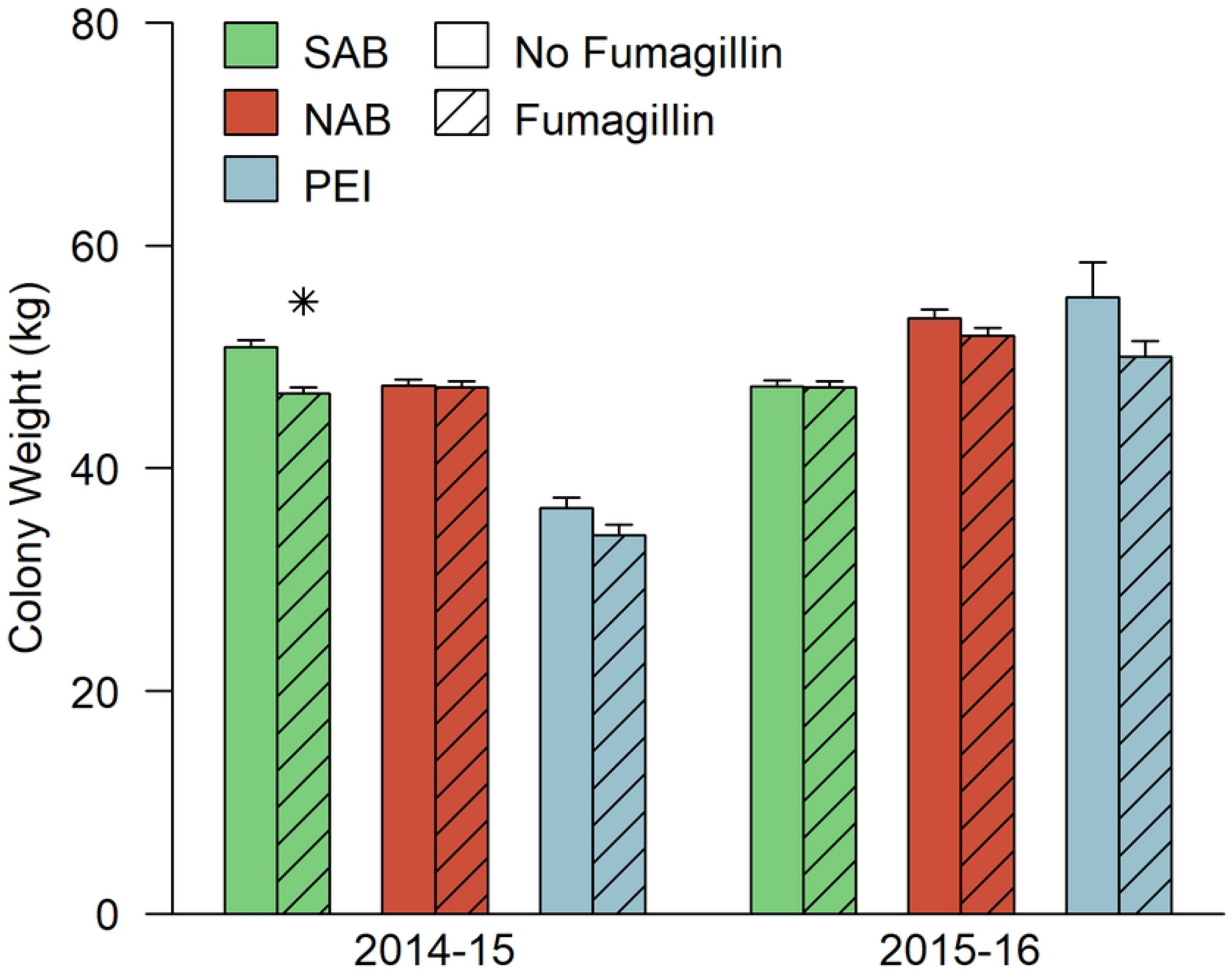
The model effect of fumagillin on colony weight (mean ± SE). Estimated marginal means are averaged across levels of protein treatment and month because there was no significant interaction between fumagillin and these factors. Each combination of region and year is shown separately because there was a significant three-way interaction of fumagillin, region, and year. Stars (*) above a column indicate that the fumagillin-treated group was statistically different from the corresponding untreated group (p<0.05, Bonferroni adjusted; see Supplement 5). SAB:Southern Alberta; NAB: Northern Alberta; PEI: Prince Edward Island.

The addition of cluster size before winter to the models for change in cluster size and change in weight improved the overall fit of both models. Larger colonies consumed more feed, and declined more in cluster size during winter, but this effect was greater in the regions with more severe winters (Northern Alberta> PEI> Southern Alberta). Following the addition of initial cluster size to the model, effects of patties and fumagillin were still detected, but appeared to be region and date-specific interactions, rather than main effects (Supplement 5).

#### Honey production

Colonies in Northern Alberta produced dramatically more honey (110 ± 3 kg per colony) than those in Southern Alberta (25 ± 1 kg per colony) in 2014 (t=23.6, df=228, p<0.001). Protein supplements did not affect honey production in 2014 (t= -0.593, df = 228, p=0.55; fumagillin had not yet been applied), but in 2015 there was a significant treatment by site interaction. In Northern Alberta, colonies that received fumagillin but not excess protein supplements produced 23.5 ± 8.4 kg more honey per colony than colonies that received neither treatment (Fig 9; t=2.82, df=177, p=0.005).

**Fig 9.**
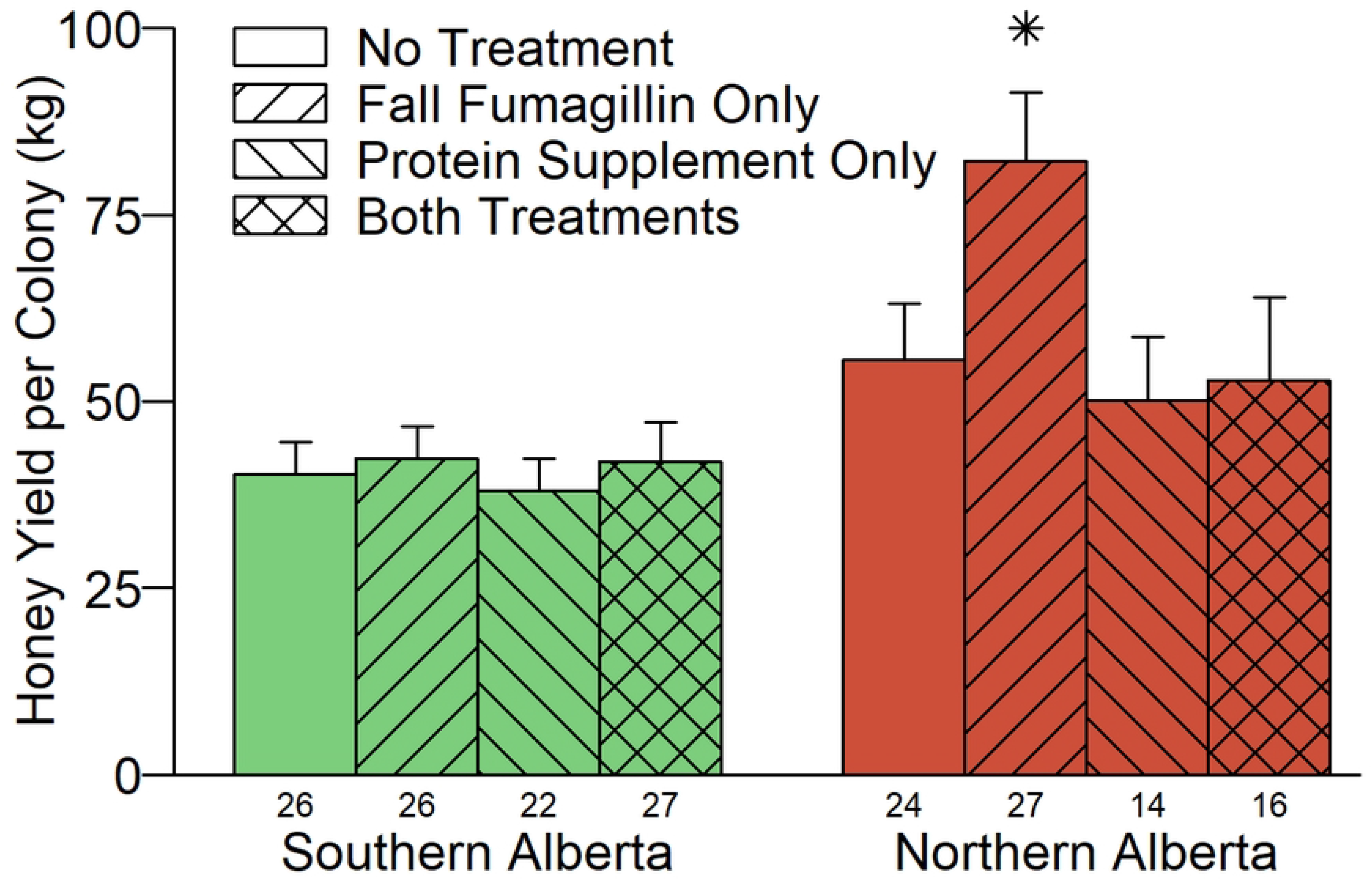
Honey yield per colony in summer 2015, by region and treatment group (mean ± SE). Stars above a column indicate that the treated group was significantly different from the untreated group (p<0.05, Bonferroni adjusted; see Supplement 5). The number of colonies that were still viable within each treatment group is shown beneath the columns. No surplus honey was produced in PEI.

#### Pollen collection

Pollen collection preceded fumagillin treatment in 2014, and fumagillin had no effect on the weight of pollen collected in 2015; as such, fumagillin treatment was dropped from the model. Since protein supplements were not applied during the bloom period of canola, all pollen collection measurements in 2014 and mid-season measurements in Alberta in 2015 measure the effect of recent, but not current, protein supplementation. The amount of pollen collected varied a great deal from day to day, likely due to local weather conditions, and also varied among colonies. In 2014, pollen was collected once from each colony, at collection dates that were distributed over the bloom period of canola. There was a significant interaction between protein supplements and region on the amount of pollen collected (F= 5.98, df=1, 223, p= 0.015). Southern Alberta colonies that had been exposed to extra protein supplements collected 2.4 ± 1.1 g less pollen per day (Fig 10; t=2.246, df=5, p=0.075) than colonies that had not been supplemented. In Northern Alberta, protein supplemented colonies collected slightly more pollen, but the difference was not significant (t=1.27, df=4, p=0.27).

**Fig 10.**
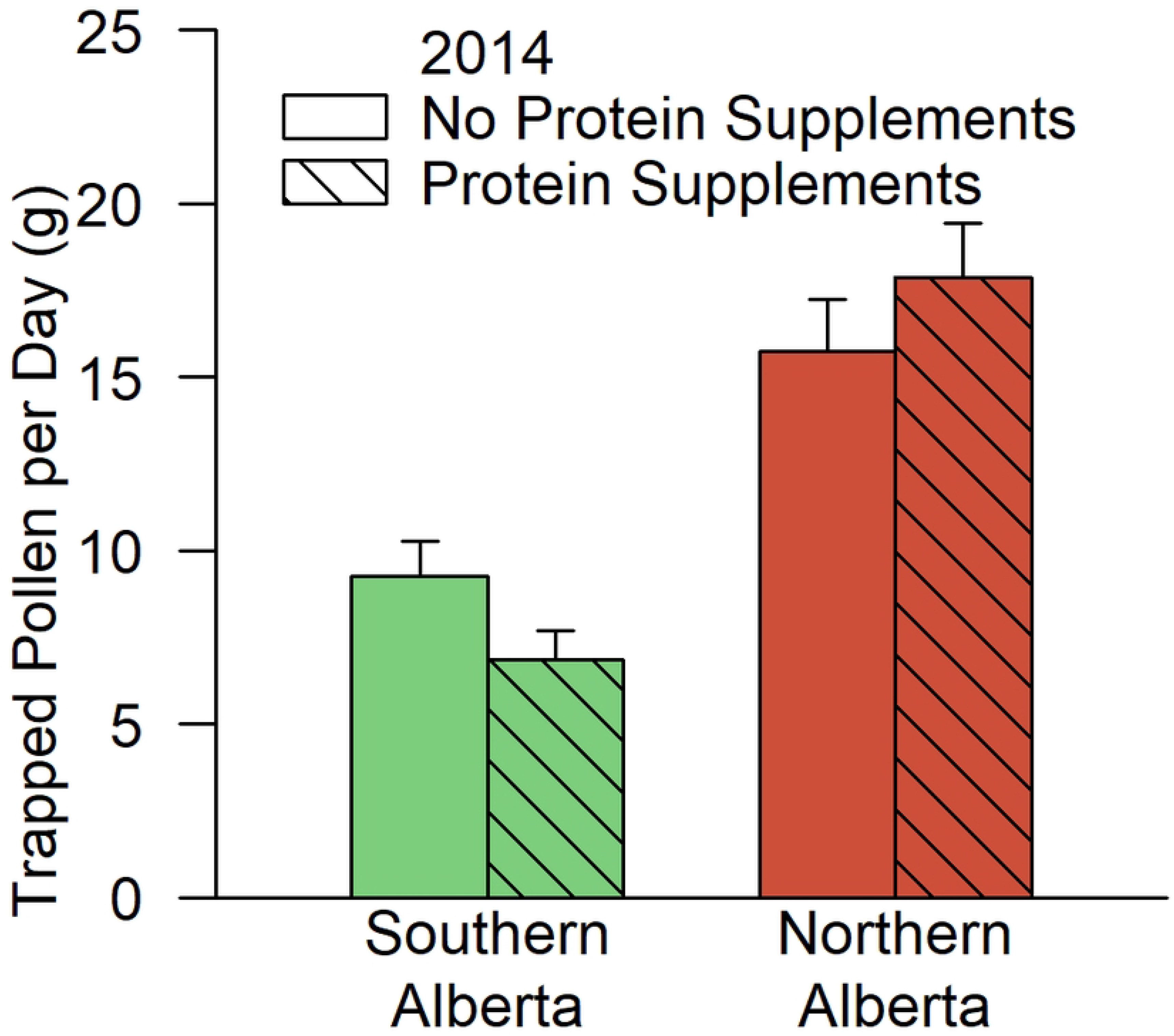
The effect of protein supplements on the quantity of trapped pollen per day in 2014 (mean ± SE). Pollen was trapped for periods of two to four days at intervals during the canola bloom; that is, between the first week of July and early August.

In 2015, pollen was trapped repeatedly from a subset of colonies in each apiary. Overall, protein supplemented colonies collected less pollen during the early and middle parts of the season, but more pollen after August 10^th^ (Fig 11; two-way interaction of protein and season: F = 3.37, df=2, 560, p=0.035; no contrasts of protein treatment within season were significant). In Southern Alberta, there was a surprising increase in pollen collection among supplemented colonies late in the year (Fig 11), but the three-way interaction of protein treatment, region, and season was not significant in the analysis of variance and was dropped from the model. There were also noteworthy differences among the regions in pollen availability (Fig 11). Large quantities of pollen were collected in Northern Alberta in the up to and including mid-summer, but after August 10th very little was collected. The pollen flow in Southern Alberta was less variable. Finally, despite the small size of colonies at PEI, the quantity of pollen collected there was greater than in Southern Alberta in the mid- and late season.

**Fig 11.**
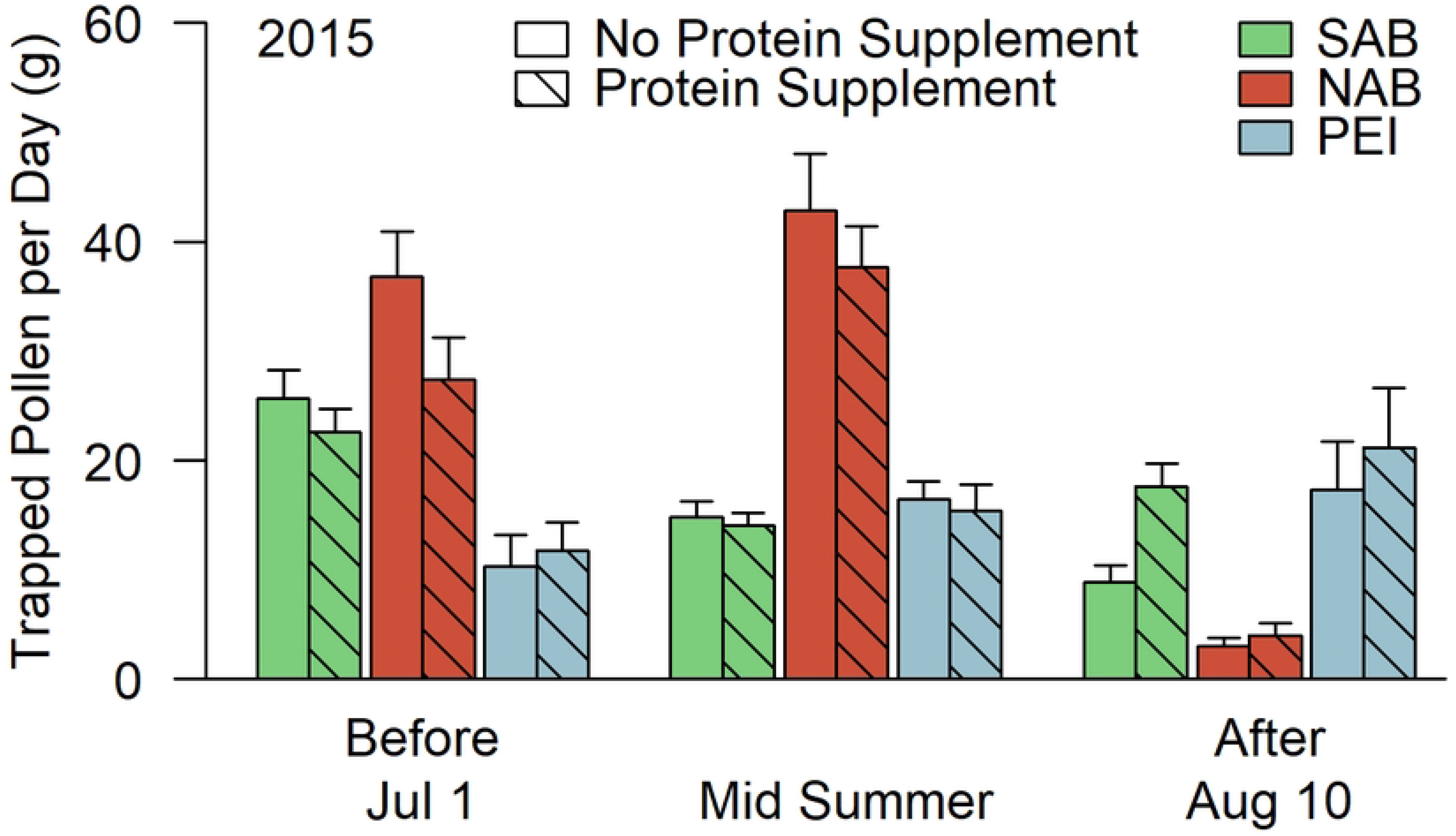
The effect of protein supplements on the quantity of trapped pollen per day in 2015 (mean ± SE). Pollen was trapped from a subset of the largest colonies in each apiary and treatment group, at two-week intervals between mid May and mid September. SAB: Southern Alberta; NAB: Northern Alberta; PEI: Prince Edward Island.

#### Colony size - an alternative view

Our analysis has treated colony survival, queen survival, and various measures of colony size and productivity as discrete outcomes, but in fact, they are likely to be related. In particular, factors that affect worker bee health, and therefore colony size and productivity, are likely also to affect the risk of queen or colony death. If a management strategy increases the size of viable colonies and also reduces the risk of colony death, either measure in isolation will underestimate the effectiveness of that strategy. Figure 12 provides a simple way to test whether this may have occurred, in which all the bees in each treatment group and region were summed on the last inspection date of the study. From this, three findings readily became apparent:

1. Local factors had a larger effect than either the fumagillin or protein supplement treatments on the population outcome.
2. Although we detected main effects related to protein supplements, the detrimental effects occurred primarily at the NAB site.
3. Although we detected only temporary and local effects from fumagillin, by the end of the study, every fumagillin-treated subgroup had considerably more bees than the corresponding group that was not fumagillin-treated.

**Fig 12.**
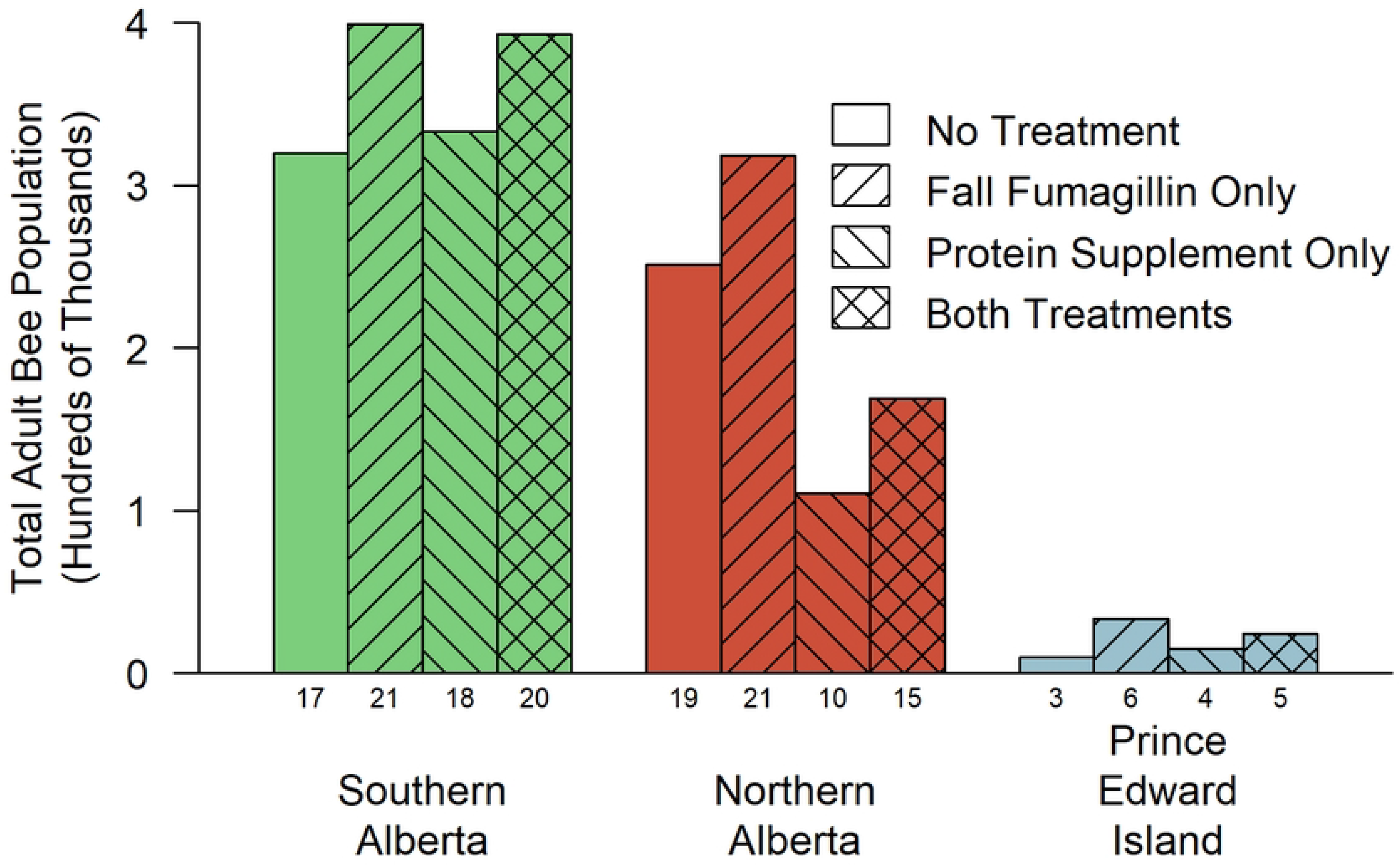
Total Adult Bee Population after Two Years of Treatment. The sums of the adult bee populations of all surviving colonies on the last inspection date of the experiment, shown by treatment group and region. Numbers beneath the columns indicate the number of viable colonies remaining.

## Discussion

In this paper we report the effects of two often-recommended beekeeper interventions on cohorts of honey bee colonies in three economically different beekeeping operations in three distinct climatic regions. The first and clearest of our observations is that the effects of treatment were small compared to the differences among the beekeeping operations, and were also small compared to the differences among colonies within an operation. We needed the full statistical power of this large, repeated-measures study to detect them. For example, one of our major findings is that protein supplements unexpectedly reduced adult bee populations over time. Yet after two years of treatment, protein supplements had reduced the average colony size in Northern Alberta by only about 1,500 bees, which was less than 1/4 of the standard deviation among colonies there at that time (6,300 bees) and just over 1/10th of the average colony size (13,000 bees). It is apparent that location, general management, and differences among individual colonies at the same site are far more influential than the two treatment strategies we tested.

### Fumagillin

Fumagillin was first reported to be effective against *Vairimorpha apis* infections in honey bees in 1952 [42], and subsequently came to be recommended for widespread prophylactic use, especially in the commercial queen industry and in the hiving of package bees [27]. Subsequently, cage trials and field trials clearly showed that fumagillin kills both species of *Vairimorpha* that infect honey bees [23, 24, 43, 44]. Despite this finding, some reviewers have recently expressed skepticism about the effectiveness of fumagillin [28, 29, 45]. Others have challenged the importance of *V. ceranae* as a pathogen [26]. Some studies have reported no significant benefits following fumagillin treatment [30, 46, 47], while others have reported large effects [48, 49]. The suspicion that fumagillin, though it kills *Vairimorpha,* may not produce reliable economic benefits is a serious concern for beekeepers.

Our results support the view that fumagillin has a positive effect on honey bee colony health. Contrary to historical recommendations [27], we did not find evidence that fumagillin significantly reduces the frequency of queen loss and the effect of fumagillin on colony survival, though positive, was also not significant. We did identify a potential negative effect of fumagillin at the colony level. Colonies accepted less feed when fumagillin was in the syrup, thus increasing the risk of starvation. Although this effect was only significant in Southern Alberta in 2014, we suspect that our data underestimate the true effect of fumagillin on feed acceptance. Colonies were provided additional unmedicated feed after the fumagillin treatment, following the beekeepers’ normal practice, and colonies were not weighed until the end of the feeding period. Weights of fumagillin-treated colonies were lower in every region-date combination except for SAB in 2015, which was also the only case where the entire fumagillin treatment was provided as a drench treatment rather than in feed.

Fumagillin did not affect brood or bee populations on most dates. However, in Northern Alberta, fumagillin was associated with higher adult bee populations in June 2015 and subsequently with dramatically higher honey production. In addition, by the end of the two-year study, fumagillin-treated groups in all regions had considerably more bees in total than the corresponding untreated groups.

### Protein supplements

Field studies have not consistently shown that commercial protein supplements are beneficial to honey bee colonies [20], although benefits have been detected in specific cases [18, 50]. Protein supplements support brood rearing when pollen availability is the limiting factor [50], but not otherwise, and many of the commercial supplements may be about equally effective [18]. However, few if any studies have tracked the same colonies for longer than one year, so the long-term effect of protein supplementation is largely unknown. Many have suggested that modern agricultural land use is a cause of malnutrition for bees [5, 51], and as such we expected that if a highly nutritious protein supplement were provided continuously, colony performance might improve.

As expected, protein supplements initially increased brood production, however, the increase was trivial in two regions and never resulted in a larger adult bee population. In one region (SAB) protein-fed colonies produced significantly more brood in June 2014 and were larger than unfed colonies in November 2014. Nevertheless, that was the last time a positive effect was observed. Across all regions, protein supplemented colonies declined more during winter than unsupplemented colonies. Over the long term, supplemented colonies were smaller than control colonies (NAB, PEI) or were not improved (SAB), and in Northern Alberta there was a substantial increase in the risk of death.

It is important to note that our protein supplement treatment was over and above the standard supplementation for the region, and our “unsupplemented” colonies received some protein supplementation in early spring, as is standard beekeeping practice. With that caveat, we draw several conclusions from these observations. First, nothing in our data supports the hypothesis that inadequate pollen (either quantity or quality) is a major cause of honey bee colony losses or under-performance in these regions. Undoubtedly, poor nutrition *could* weaken or kill hives, but in regions with a successful bee industry the pollen supply is probably adequate, and not the cause of recent poor performance. Secondly, protein supplements increase the rate of brood rearing at certain times and places - notably Southern Alberta in the first year. Thirdly, protein supplements consistently led to lower adult bee populations. This suggests that the increased brood rearing may have been offset by a decrease in the average adult lifespan, and the harmful effect on the adults was more significant over the long term than the benefit to the brood.

We are not the first to suggest that protein supplements lead to shorter lifespans for adult bees. La Montaigne et al. [52] fed two protein supplements to colonies in Quebec and reported no improvement in colony performance as a result. They examined the effect on adult bee lifespan using marked bees in the hive and found that both supplements reduced average adult lifespans. Cage trials have shown that when *Vairimorpha*-infected bees consume protein, two effects occur: (1) infected bees that receive protein supplements live longer than infected bees that did not receive protein supplements and (2) *Vairimorpha* spores replicate faster and to a higher maximum level in the midguts of these bees [53, 54]. It has been unclear how these results relate to colony level performance, but the following interpretation would be consistent with our observations: when an infected bee consumes protein, both the bee and the parasite benefit, however the benefit to the bee is limited to the life of that bee, while the additional parasites remain in the hive to increase the prevalence of the disease. During this study, we collected over 6000 samples of bees for analysis of pests and parasites, and from analyses of these samples, we intend to address this question in a future report.

The value of high quality, nutritious natural pollen for bee hives has been established beyond any possible doubt [55]. In this study, the patties contained 25% pollen, which is far more than most commercial formulations, but they did not produce the expected benefits. Four possible explanations occur to us: (1) as mentioned, the colonies may not have been pollen-limited; (2) there might be some anti-nutritional compound or pesticide, unknown to us, remaining in the formulation; (3) periods of reduced pollen consumption (and hence reduced brood rearing) might be required for natural disease and pest resistance in the honey bee; or (4) the unnatural way in which protein supplements are supplied to the hive may have unintended consequences. Pollen that is naturally collected by the bees is stored as bee bread and is consumed by both nurse bees and larvae [56]. Noordyke et al. [57] recently examined the fate of protein supplements fed as patties in the hive, and found that the supplements were not stored as bee bread or fed directly to larvae; they were apparently consumed almost entirely by the adult bees.

### Regional differences

Hives belonging to the Southern Alberta seed canola pollinator had by far the greatest survival (73% survived two years) and the most stable colony populations. The hives grew slowly during summer but declined only slightly in winter. These large and stable populations are beneficial in a pollination context. However, only modest amounts of honey and pollen were produced. In the first year these bees benefited from the extra protein supplement, which suggests their environment was slightly pollen-limited in spring and possibly fall 2014, but since the difference was small and transient, it was unlikely to have justified the cost of treatment.

Hives from the Northern Alberta honey producer, which was the bee research laboratory of AAFC’s Beaverlodge Research Farm, produced far more honey and pollen than those from SAB, especially in the first year, as is typical of colonies in this region [58]. Colonies in NAB grew much faster each summer, but also declined much more during winter. The quantity of trapped pollen dropped to near zero after the first week of August, and remained low; brood rearing declined more slowly, but was predominantly complete by the end of September. These colonies faced the longest winter of the three sites, and depended greatly on the longevity of their winter bees.

The poorest performing hives in the study were those of the PEI blueberry pollinator, nearly all of which died over the two-year study. The poor performance of these hives, in part, may have been due to the study protocol. We required nucleus colonies to be started as early in the year as possible in order to match the treatments used in Alberta. However, the cooperating producer had a regular practice of splitting colonies in August, after blueberry pollination, and carried out that practice on the experimental colonies which had already been split. Thus, even though these colonies had the highest rate of brood production observed in the study, they were extremely small in fall and the splits had a higher-than-average rate of death. Nevertheless, the study protocol was likely only an incidental contributor to colony mortality in the region. Also noteworthy was the larger number and variety of visible disease symptoms in these colonies than anywhere else, higher varroa mite infestations, and during the first year of the study, colonies that were inadequately fed prior to winter resulting in high rates of starvation.

## Conclusions

We have examined the long-term effects of two beekeeper interventions on honey bee colony health. Protein supplementation and fumagillin treatment both produced benefits in specific situations, but not in general. Local and individual differences among colonies were far larger than the effects of either treatment. There appears to be no risk associated with ordinary fumagillin treatment, provided the hive has adequate feed, and in some circumstances, there are substantial benefits. It seems reasonable to expect that the benefits of fumagillin are greatest when infections with *Vairimorpha* spores are severe, which may be more likely in with regions with a long, cold, temperate winter. Beekeepers should be cautioned against applying protein supplements when there is not a clear protein shortage. It may not be economically beneficial and under certain circumstances, the supplement may lead to unintended harm.

## Acknowledgments

We thank the participation of Landen Stronks and Gabriel Calixte, of Kiwi Brian*’*s Honey, as well as Peter Dillon and Jasper Wyman and Son Canada, Inc. We also acknowledge Sean Murray for his assistance in the conduct of the experiment in PEI. Thanks also goes to technical assistance from: Elena Battle, Jamie Malbeuf, Marika Viens, Zachary Wagman, Benjamin King, Breanalee Beer, Ian Johnson, Danielle Ediger, Justin Mufford, and Christopher Isaac. This project was supported by AAFC through Project # J-000049 *“*Health of Bee Pollinators in Canadian Agriculture*”*.

## Supporting information

**S1 Methods Details.** Includes Tables S1 - S8 and Fig S1, referred to in the text.

**S2 Main Dataset.** Spreadsheet in .csv format.

**S3 Pollen 2015 Dataset.** Spreadsheet in .csv format.

**S4 Statistical Analysis Code.** R markdown file with the statistical analysis code.

**S5 Statistical analysis output file.** HTML file showing the output of the statistical analysis.

